# Genomic Characterization of a Typical Enteropathogenic *Escherichia coli* Isolate from an Egyptian Fruit Bat, *Rousettus aegyptiacus* from Coastal Kenya

**DOI:** 10.64898/2026.11.19.767828

**Authors:** Jacob Ngugi Kamau, Bryson Kimemia, David Abuom, Benard Agwanda, Lillian Musila, Erick Odoyo, Mary W. Kivata, Cecilia K. Mbae, Justus Onguso, Haynes Robert, Gerald G. Kellar, Joel Lutomiah, John Eads, Fredrick Eyase

**Affiliations:** Department of Emerging Infectious Diseases, Walter Reed Army Institute of Research-Africa, Kenya; Centre for Virus Research, Kenya Medical Research Institute, Nairobi, Kenya; Center for Microbiology Research, Kenya Medical Research Institute, Nairobi, Kenya; Centre for Biotechnology Research and Development, Kenya Medical Research Institute, Nairobi, Kenya; Institute for Biotechnology Research, Jomo Kenyatta University of Agriculture and Technology (JKUAT), Nairobi, Kenya; National Museums of Kenya, Nairobi; Department of Biological and Physical Sciences, Karatina University, Karatina, Kenya

**Keywords:** *Escherichia coli*, enteropathogenic *Escherichia coli* (EPEC), whole-genome sequencing, Egyptian fruit bat, *Rousettus aegyptiacus*, wildlife genomics, One Health, virulence factors, antimicrobial resistance, Kenya

## Abstract

*Enteropathogenic Escherichia coli* (EPEC) is an important diarrheagenic pathogen responsible for substantial morbidity among young children, particularly in low- and middle-income countries. Although atypical EPEC has frequently been reported in animal hosts, information on the occurrence and genomic characteristics of typical EPEC in wildlife reservoirs remains limited. This study characterized *Escherichia coli* isolated from an Egyptian fruit bat (*Rousettus aegyptiacus*) sampled in coastal Kenya using whole-genome sequencing to evaluate virulence determinants, antimicrobial resistance genes, and phylogenetic relationships relevant to public health. Oral and rectal swabs were collected from 24 bats captured in Kikambala, Kilifi County, Kenya. Bacterial isolates were cultured, sequenced on the Illumina NextSeq 1000 platform, and analyzed using a whole-genome sequencing workflow that included taxonomic classification, *in silico* serotyping and multilocus sequence typing, virulence gene profiling, antimicrobial resistance gene detection, plasmid identification, and core genome phylogenetic analysis. Among 18 *E. coli* isolates identified, representing 13 serotypes, one isolate, KIKB-471-SWR-A (O128:H6; ST15625), was classified as typical EPEC. Genomic analysis demonstrated the presence of the chromosomal locus of enterocyte effacement (LEE), including the *eae* and *tir* genes, together with a complete plasmid-associated bundle-forming pilus (*bfp*) operon, confirming the characteristic virulence architecture of typical EPEC. The isolate also encoded a complete type III secretion system and multiple additional adherence, secretion, and iron-acquisition determinants. Although numerous antimicrobial resistance-associated genes, including multidrug efflux systems and chromosomal β-lactamase genes, were detected, the isolate remained phenotypically susceptible to all antimicrobials tested. Core genome phylogenetic analysis placed the isolate within the EPEC lineage, clustering most closely with a human-derived EPEC strain 1554 while remaining genetically distinct from both commensal and laboratory *E. coli* reference strains. The identification of a genomically confirmed typical EPEC isolate in a free-ranging Egyptian fruit bat expands current knowledge of the diversity of EPEC circulating in wildlife. These findings demonstrate that wildlife can harbor *E. coli* lineages possessing virulence determinants associated with human pathogenic EPEC and underscore the importance of genomic surveillance at wildlife–human interfaces within a One Health framework. Although this study does not demonstrate zoonotic transmission, it provides genomic evidence supporting continued surveillance of wildlife-associated *E. coli* to improve understanding of the ecology, evolution, and public health significance of diarrheagenic *E. coli*.

## Background

*Escherichia coli* is a genetically diverse bacterial species that occurs predominantly as a commensal inhabitant of the gastrointestinal tract of humans and animals. However, acquisition of virulence-associated genetic elements through horizontal gene transfer has given rise to pathogenic lineages capable of causing both intestinal and extraintestinal disease (1–8). Diarrheagenic *E. coli* (DEC) comprise several pathotypes distinguished by their virulence mechanisms, including enteropathogenic *E. coli* (EPEC), enterohemorrhagic (EHEC), enterotoxigenic (ETEC), enteroaggregative (EAEC), enteroinvasive (EIEC), and diffusely adherent *E. coli* (DAEC) (4,8). Collectively, DEC remain among the leading bacterial causes of childhood diarrheal disease in low- and middle-income countries, contributing substantially to morbidity and mortality among children younger than five years (9,10).

EPEC is characterized by the formation of attaching-and-effacing (A/E) lesions on intestinal epithelial cells, mediated by the chromosomal locus of enterocyte effacement (LEE). The LEE encodes a type III secretion system (T3SS), its associated translocated effector proteins, the adhesin Intimin (encoded by *eae*), and the translocated Intimin receptor (*tir*), which together facilitate intimate bacterial attachment and cytoskeletal remodeling of host cells (2,8,11). Typical EPEC (tEPEC) additionally possesses the EPEC adherence factor (EAF) plasmid encoding the bundle-forming pilus (BFP), a type IV fimbria that mediates localized adherence during the early stages of colonization (12–14). In contrast, atypical EPEC (aEPEC) lacks the EAF plasmid and is frequently recovered from animal hosts while exhibiting greater genomic diversity (11,14,17,18).

The emergence and dissemination of EPEC lineages are driven by the acquisition, maintenance, and loss of mobile genetic elements carrying virulence and antimicrobial resistance determinants. Whole-genome sequencing (WGS) has substantially improved the resolution with which these evolutionary processes can be investigated by enabling simultaneous characterization of serotypes, sequence types, phylogenetic relationships, virulence gene repertoires, plasmid content, and antimicrobial resistance determinants. Consequently, WGS has become an important tool for investigating the ecology and evolution of pathogenic *E. coli* within a One Health framework that recognizes the interconnectedness of human, animal, and environmental health.

Bats represent one of the most species-rich mammalian orders, and occupy diverse ecological niches across tropical and subtropical ecosystems (19,20). Their long lifespan, colonial roosting behavior, extensive movement, and frequent interactions with livestock, domestic animals, and humans have attracted increasing attention regarding their role in the maintenance and dissemination of microorganisms of public health and veterinary importance (21–26). Although bats are well recognized as reservoirs of numerous viral pathogens, comparatively less is known about the genomic diversity and pathogenic potential of bat-associated enteric bacteria, particularly diarrheagenic *E. coli*. Most available studies have relied on PCR-based detection of virulence genes, whereas comparatively few have employed whole-genome sequencing to comprehensively characterize bat-derived EPEC isolates.

In Kenya, information regarding the genomic characteristics of wildlife-associated *E. coli* remains limited despite increasing human encroachment into wildlife habitats and growing recognition of the importance of wildlife surveillance. Characterizing virulence determinants, antimicrobial resistance genes, and evolutionary relationships among bat-associated *E. coli* can improve understanding of the ecology of pathogenic lineages and inform surveillance programs aimed at identifying microorganisms with potential public health relevance. In this study, we performed whole-genome sequencing and comparative genomic analysis of *E. coli* isolated from an Egyptian fruit bat (*Rousettus aegyptiacus*) sampled in coastal Kenya. We characterized serotypes, sequence types, virulence gene repertoires, antimicrobial resistance determinants, plasmid content, and phylogenetic relationships, with particular emphasis on the genomic characterization of a typical EPEC isolate and its implications for public health surveillance within a One Health framework.

## Materials and Methods

### Field Site and Sample Collection

Kikambala, located in Kilifi County, is a coastal area with an altitude of between 60 – 150m above sea level. The county is humid with a bimodal rainfall pattern, temperatures ranging from 21 - 30ºC annually, with vegetation predominantly tropical flora. Kilifi has caves both within and outside of the Arabuko-Sokoke forest ecosystem.

Bats were captured using 12m mist nets by a team of qualified personnel employing the use of personal protective equipment. The sampling was done in December, 2024, with selection of sites based on information about observed colonies and roosts in Kikambala. The phenotypic identification of the captured bat species was conducted using taxonomic field keys, with oral and rectal swabs then collected and stored in AMIES transport media. The labeled oral and rectal swabs, were put into zip lock bags and immediately deposited in dry ice in a cooler box for transportation. Once in the laboratory, they were stored at -80 °C prior to further analysis. Due to threats on their decreasing population, and their ecological importance, all captured bats were released after collection of the samples.

### Bacteriological Cultures and Antimicrobial Susceptibility Testing

The swabs were retrieved and thawed at room temperature before being inoculated on 5% Sheep Blood Agar (SBA) and MacConkey Agar (MAC); and incubated at 37ºC for 24 to 48 hours aerobically. Sub-cultures were undertaken on both SBA and MAC to obtain pure bacterial isolates morphologically. The antimicrobial susceptibility testing (AST) was performed on fresh pure bacterial cultures, using the VITEK 2 AST-GN83 (BioMérieux, La Balme les Grottes, France) card, and interpreted based on the Clinical and Laboratory Standards Institute (CLSI) guidelines (27).

### DNA Extraction, Library Preparation and Sequencing

DNA was extracted from the pure *E. coli* isolates using the Zymo Quick-DNA Fungal/Bacterial Miniprep Kit (Orange, California) per the manufacturer’s instructions (28), and the concentration quantified using the Qubit dsDNA High Sensitivity Assay Kit (Thermo Fisher Scientific, Waltham, USA). For whole-genome sequencing (WGS), multiplexed DNA libraries were prepared using the DNA Prep Kit (Illumina Inc., San Diego, USA), and subsequently sequenced on the Illumina NextSeq 1000 System, using a NextSeq P1 Reagent Kit (300 cycles) to obtain a 2 x 150 bp paired-end read output.

### Quality Control and Assembly

Demultiplexed paired-end reads were evaluated for quality using FastQC v0.12.1. Adapter sequences and low-quality bases were removed using Trimmomatic v0.40, with *de novo* assembly of bacterial genomes performed using Unicycler v0.5.1. The assembly quality was assessed with QUAST v5.3.0. Final assemblies were retained if they contained contigs >500 bp in length and exhibited >5× k-mer coverage.

### Taxonomy identification, *In Silico* Serotyping, and Sequence Type Determination

Taxonomic classification of isolates was performed using Kraken2 v2.1.3 to confirm the bacterial species. *In silico* serotyping was conducted on ABRicate v1.0.1, employing the EcOH database. Sequence typing was carried out using the MLST (v2.23.0) tool (Achtman 7 database), with confirmation of sequence types and the assignment of phylogroups undertaken on the EnteroBase webserver (https://enterobase.warwick.ac.uk/).

### Virulence and Antimicrobial Resistance Gene Detection

Contigs were screened for virulence-associated genes on ABRicate v1.0.1 employing the Virulence Factor Database (VFDB), with antimicrobial resistance determinants detected using the Comprehensive Antimicrobial Resistance Database (CARD), and the ResFinder database. Resistance plasmids were identified through the PlasmidFinder database. Virulence gene presence was determined based on thresholds of >90% coverage and >95% sequence identity.

### Phylogenetic Tree Construction

Comparative analysis included the reference EPEC strain E2348/69, the human clinical EPEC strain 1554, the food-associated EPEC strain 1668-4a, the laboratory strain K-12 MG1655, and the commensal strain HS. These genomes represent well-characterized reference strains encompassing clinically relevant EPEC lineages, non-pathogenic laboratory and commensal lineages, and epidemiologically distinct isolates, thereby providing a robust phylogenetic framework for interpreting the bat-derived isolate. Core genome alignment was performed using ParSNP v2.1.4, with *Escherichia coli* EPEC strain E2348/69 (GCF_000026545.1) designated as the reference genome. Regions of recombination were excluded from the resulting core single nucleotide polymorphism (SNP) alignment. A maximum-likelihood phylogeny was inferred with RAxML v1.2.2, employing the GTRGAMMA substitution model and 1000 bootstrap replicates to assess branch support. Phylogenetic trees were visualized and edited using iTOL (version) (https://itol.embl.de/).

## RESULTS

### Taxonomy identification, Serotypes, and Sequence Types

A total of 48 oral and rectal swabs were collected from 24 bats in Kilifi County, predominantly *Rousettus aegyptiacus* (n=23), with one bat identified as *Taphozous hildegardeae*. Following subculture, all 67 pure bacterial isolates obtained were taxonomically classified. Among these, 18 isolates were identified as *Escherichia coli*, comprising 13 distinct O:H serotypes (29,30). The most frequently observed serotypes were O107:H45 and O112:H1 (each n=3), followed by O8:H7 (n=2), while the remaining serotypes had single isolates. *E. coli* isolate, KIKB-471-SWR-A, recovered from a bat rectal swab was taxonomically identified as enteropathogenic *E. coli* (EPEC O128:H6), having a sequence type, ST15625, and belonging to phylogroup B2 (31,32).

### Virulence Gene Profiles

Comparative genomic analysis of EPEC isolate O128:H6, with unclassified pathotypes O88:H5 and O83:H7, revealed overlapping but distinct virulence repertoires (Table 1). The genome for isolate O128:H6 contained the *eae* gene encoding Intimin, and the complete *bfp* operon (*bfpA–bfpO*). Genes encoding the type III secretion system (T3SS) were present, including structural components (*escC– escV*), chaperones (*cesAB, cesD, cesD2, cesL, cesT*), and regulators (*sepD, sepL*). The *tir* gene and non-LEE effector *nleF* were also detected in the isolate supporting conserved attaching-and-effacing mechanisms. Effector diversity was also evident, O128:H6 carried *espB, espD, espG* genes, whereas O83:H7 harbored *espR1, espX4, espX5*, suggesting functional divergence in host interaction strategies. Adhesin-related genes common across the isolates included type I fimbriae and the *E. coli* common pilus (ECP). Notably, O128:H6 lacked *fimA and fimB*, potentially impairing pilus assembly and phase variation, while O83:H7 retained the full *fim* cluster minus *fimA*. Iron acquisition systems were also strongly represented across all isolates, including genes encoding ferric enterobactin uptake, heme uptake (*chuA–chuY*), and yersiniabactin siderophores (*fyuA, irp1, irp2, ybtA–ybtX*). The *astA* gene was detected in isolate O128:H6, but annotation revealed it encodes arginine N-succinyltransferase, and not the heat-stable enterotoxin EAST1 associated with enteroaggregative *E. coli* (EAEC).

**Table 1.** Comparative distribution of virulence genes among *E. coli* isolates O88:H5, O128:H6, and O83:H7. Genes are grouped by functional categories: adhesion, toxin production, secretion systems, effector proteins, iron acquisition, and intimin/attachment. The O128:H6 isolate displayed the complete *bfp* operon, the Intimin gene, and the T3SS secretion system gene repertoire, while O83:H7 exhibited unique effector diversity. Shared systems included type 1 fimbriae genes, E. coli common pilus (ECP) genes, general secretion pathway (type II secretion system) genes, and ferric enterobactin uptake and processing genes.

| **1.Adhesion** | **O88:H5** | **O128:H6** | **O83:H7** |
|---|---|---|---|
| Bundle-forming pilus (BFP) genes |  | bfpA, bfpB, bfpC, bfpD, bfpE, bfpF, bfpG, bfpH, bfpI, bfpJ, bfpK, bfpL, bfpP, bfpU |  |
| Type 1 fimbriae genes | fimF, fimG, fimH | fimC, fimD, fimE, fimF, fimG, fimH, fimI | fimB, fimC, fimD, fimE, fimF, fimG, fimH, fimI |
| Associated with F1C fimbriae, | focH |  |  |
| P fimbriae genes | papC, papD, papE, papH, papJ, papK |  |  |
| S fimbriae genes | sfaB, sfaD, sfaE, sfaF, sfaG, sfaX, sfaY |  |  |
| E. coli common pilus (ECP) genes | yagV/ecpE, yagW/ecpD, yagX/ecpC, yagY/ecpB, yagZ/ecpA, ykgK/ecpR | yagV/ecpE, yagW/ecpD, yagX/ecpC, yagY/ecpB, yagZ/ecpA, ykgK/ecpR | yagV/ecpE, yagW/ecpD, yagX/ecpC, yagY/ecpB, yagZ/ecpA, ykgK/ecpR |
| **2.Toxin Production** |  |  |  |
| Subtilase-like toxin | aslA | aslA |  |
| Hemolysin | hlyB, hlyD |  |  |
| Enterobactin biosynthesis and transport genes | entA, entB, entC, entD, entE, entF, entS | entA, entB, entC, entD, entE, entF, entS |  |
| **3.Secretion Systems** |  |  |  |
| Chaperone and regulatory proteins for the T3SS |  | cesAB, cesD, cesD2, cesL, cesT |  |
| Structural and functional components of the T3SS |  | escC, escD, escE, escF, escG, escI, escJ, escL, escN, escO, escR, escS, escT, escU, escV |  |
| T3SS-associated genes |  | sepD, sepL, sepQ/escQ |  |
| General secretion pathway (type II secretion system) genes | gspC, gspD, gspE, gspF, gspG, gspH, gspI, gspJ, gspL, gspM | gspC, gspD, gspE, gspF, gspG, gspH, gspI, gspJ, gspK, gspL, gspM | gspC, gspD, gspE, gspF, gspG, gspH, gspI, gspJ, gspK, gspL, gspM |
| **4.Effector Proteins** |  |  |  |
| T3SS effector proteins |  | espB, espD, espG | espR1, espX4, espX5 |
| Non-LEE-encoded effector |  | nleF |  |
| Translocated intimin receptor |  | tir |  |
| **5.Iron Acquisition** |  |  |  |
| Heme uptake system genes | chuA, chuS, chuT, chuU, chuV, chuW, chuX, chuY | chuA, chuS, chuT, chuU, chuV, chuW, chuX, chuY |  |
| Ferric enterobactin uptake and processing genes | fepA, fepB, fepC, fepD, fepG, fes | fepA, fepB, fepC, fepD, fepG, fes | fepA, fepB, fepC, fepD, fepG, fes |
| Salmochelin siderophore system genes | iroB, iroC, iroD, iroE, iroN |  |  |
| Yersiniabactin siderophore system genes | ybtA, ybtE, ybtP, ybtQ, ybtS, ybtT, ybtU, ybtX |  |  |
| Additional yersiniabactin-related genes | fyuA, irp1, irp2 |  |  |
| **6.Intimin and Attachment** |  |  |  |
| Intimin |  | eae |  |
| Intimin-like protein |  | etgA |  |
| **7.Outer Membrane Proteins** |  |  |  |
| Outer membrane protein | ompA | ompA | ompA |
| Adhesion and colonization factor | fdeC | fdeC | fdeC |

### Phenotypic Antimicrobial Susceptibility and Resistance Genomic Profile

EPEC isolate (O128:H6) was susceptible to all tested antibiotics (Ampicillin, Amoxicillin/Clavulanic Acid, Ampicillin/Sulbactam, Piperacillin/Tazobactam, Cefazolin, Cefuroxime, Cefurotaxime Axetil, Cefoxitin, Cefotaxime, Ceftazidime, Ceftriaxone, Cefepime, Aztreonam, Meropenem, Amikacin, Gentamicin, Ciprofloxacin, Nitrofurantoin, Trimethoprim/Sulfamethoxazole). Genomic profiling of the isolate revealed a diverse set of antimicrobial resistance (AMR) determinants, primarily associated with multidrug efflux systems and β-lactamase activity. The *acrA* and *acrB* genes, together with *TolC*, constituted the AcrAB-TolC efflux complex (33), conferring resistance to multiple antibiotic classes including cephalosporins, fluoroquinolones, glycylcyclines, penams, phenicols, rifamycins, tetracyclines, and triclosan. Additional efflux transporters were identified, including AcrD, regulated by *baeRS* and *cpxAR (33)*, which mediates aminoglycoside resistance, and *acrE/acrF*, linked to cephalosporin, cephamycin, fluoroquinolone, and penam resistance. Multiple Mdt family proteins (MdtA–MdtP) were detected, forming complexes such as MdtABC-TolC (33), associated with resistance to aminocoumarins, fluoroquinolones, macrolides, penams, fosfomycin, puromycin, and acriflavine. The *bacA* gene (34), involved in cell wall biosynthesis, conferred resistance to bacitracin. Genes associated with resistance to antimicrobial peptides were also present, including *ugd, eptA*, and *pmrF*, which mediate polymyxin resistance (35), and *yojI*, which forms the YojI-TolC complex to expel microcin molecules (36). Additional efflux systems included EmrA, EmrB, and EmrK, forming complexes with TolC and conferring resistance to fluoroquinolones and tetracyclines. Mutations in *emrR*, the negative regulator of EmrAB-TolC (37), are known to result in overexpression of this system. The global regulator *marA* was also detected, which induces AcrAB (33) expression under antibiotic stress, thereby broadening resistance to carbapenems, cephalosporins, cephamycin, fluoroquinolones, glycylcyclines, monobactams, penams, penems, phenicols, rifamycins, tetracyclines, and triclosan.

Other transporters included MdfA (37), linked to tetracycline resistance, and MsbA, an ATP-binding cassette (ABC) (36) transporter conferring nitroimidazole resistance. Two class C β-lactamase genes, *ampC* and *ampH*, were detected, conferring resistance to cephalosporins and penams (38). Regulatory elements such as H-NS (39) were also present, known to repress efflux pump genes (*acrE, mdtE, emrK*) and modulate fimbrial recombinases (*fimB, fimE*) involved in biofilm formation (40). Finally, plasmid replicons Col156_1, Col(BS512)_1, and ColRNAI_1 were identified, suggesting potential horizontal transfer of resistance determinants.

### Phylogenetic analysis

Maximum-likelihood phylogenetic reconstruction based on core genome SNP alignment revealed two major clades among the bat-derived *Escherichia coli* isolates (Figure 1). One clade clustered around the enteropathogenic *E. coli* (EPEC) reference genome E2348/69 (GCF_000026545.1) (41), while the second grouped with the human commensal strain HS (GCF_000017765.1) and the non-pathogenic laboratory strain K-12 (GCF_047038235.1) (6). The inclusion of HS (human commensal) provided a benchmark for assessing evolutionary divergence from typical commensal lineages (6), whereas EDL933 (GCF_029590495.1), an enterohemorrhagic *E. coli* (EHEC) O157:H7 strain (42), served as the outgroup. The bat-derived isolate *E. coli* O128:H6 clustered within the EPEC-associated clade, showing close phylogenetic proximity to EPEC 1554 (GCF_001742645.1), a human pathogenic O157:H39 strain isolated from a diarrheal case in the Netherlands. It was more distantly related to EPEC 1668-4a (GCF_021504115.1), an environmental strain recovered from beef in Australia. Other bat-derived isolates, including O88:H5, O150:H6, O150:H45, and O107:H45, also grouped within the EPEC clade, suggesting shared evolutionary ancestry and potential conservation of virulence traits

**Figure 1.**
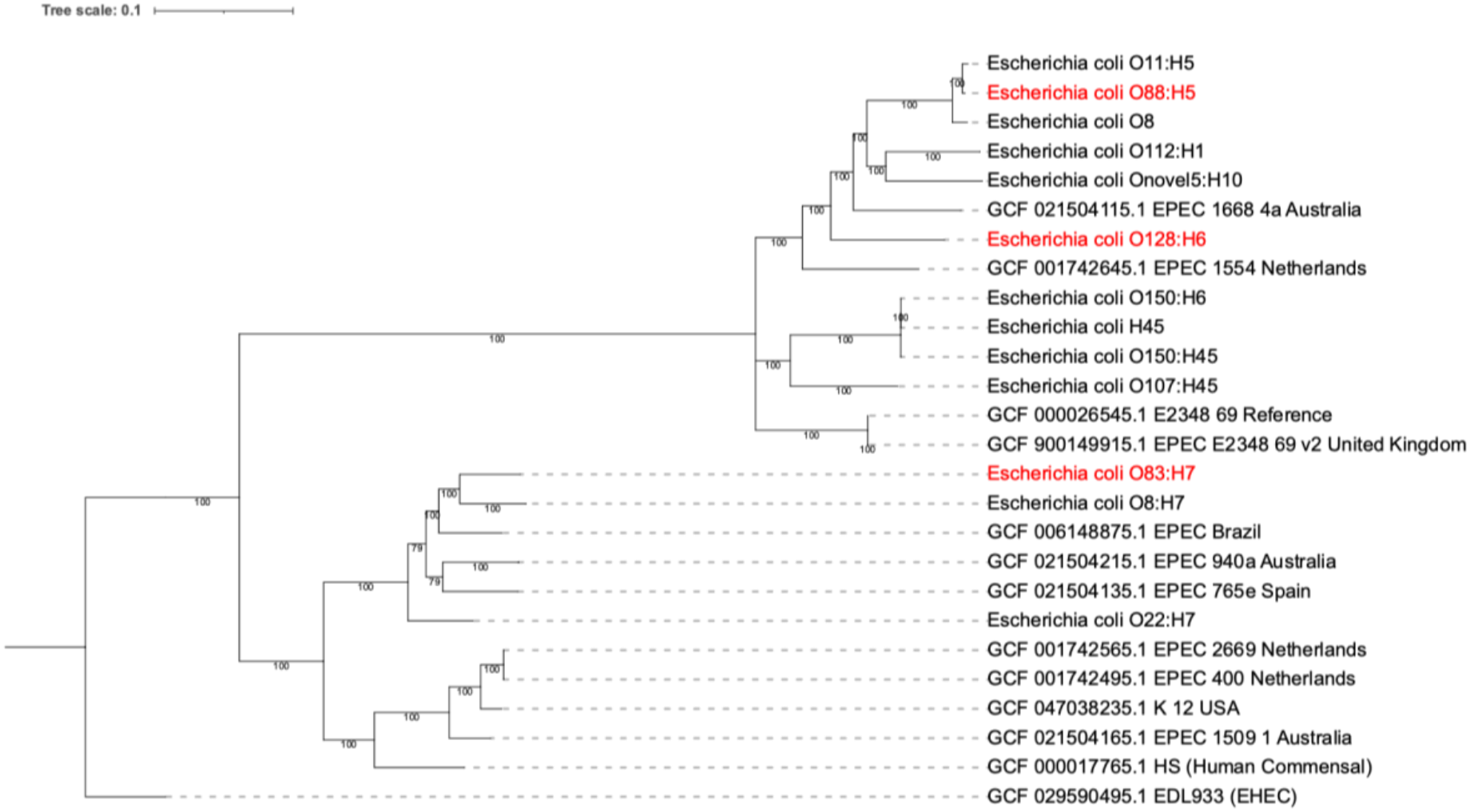
Maximum-likelihood phylogenetic tree of *E. coli* isolates based on core genome SNPs. The taxonomically identified EPEC isolate, O128:H6, and two other *E. coli* isolates are highlighted in red. Reference genomes include EPEC E2348/69, human commensal HS, laboratory strain K-12, and EHEC EDL933 (outgroup). Bootstrap values indicate branch support. Scale bar represents genetic distance.

## Discussion

This study provides a whole-genome characterization of a typical enteropathogenic *Escherichia coli* (EPEC) isolate recovered from a free-ranging Egyptian fruit bat (*Rousettus aegyptiacus*) in coastal Kenya. Among the 18 *E. coli* isolates identified, the KIKB-471-SWR-A isolate possessed the defining genomic features of typical EPEC, including the chromosomal locus of enterocyte effacement (LEE) and the plasmid-associated bundle-forming pilus (BFP) operon. Comparative genomic analysis further demonstrated conservation of the type III secretion system (T3SS), multiple adherence-associated determinants, iron acquisition systems, and close phylogenetic relatedness to established human EPEC lineages. Collectively, these findings expand current knowledge of the genomic diversity of wildlife-associated EPEC and highlight the value of whole-genome sequencing for surveillance of diarrheagenic *E. coli* within a One Health framework.

The simultaneous detection of *eae, tir*, and the complete *bfp* operon provides strong genomic evidence that isolate O128:H6 represents a typical EPEC strain. The distinction between typical and atypical EPEC is epidemiologically important because the presence of the EPEC adherence factor (EAF) plasmid encoding bundle-forming pili has historically been associated with efficient localized adherence and enhanced intestinal colonization (11–14,17). The complete complement of LEE-associated structural genes, secretion components, regulatory proteins, and translocated effectors identified in the present isolate is consistent with the conserved molecular machinery required for attaching-and-effacing lesion formation described in human EPEC (2,8,11,13,54). Although this study did not evaluate gene expression or epithelial cell adherence experimentally, the genomic architecture closely resembles that reported for well-characterized clinical EPEC strains.

The isolate also possessed several accessory virulence determinants that may enhance colonization and persistence. Conserved type 1 fimbrial genes, *E. coli* common pilus genes, and multiple iron acquisition systems, including enterobactin, heme uptake, and yersiniabactin-associated loci, were identified. Iron acquisition systems are widely recognized as important fitness determinants that facilitate bacterial survival in iron-limited host environments and frequently occur among pathogenic *E. coli* lineages (6,7). The coexistence of multiple adherence mechanisms and siderophore systems illustrates the genomic complexity of this isolate and suggests adaptation to diverse ecological niches. Nevertheless, functional studies are required to determine the contribution of these determinants to colonization or persistence in bat hosts.

An important observation was the detection of numerous antimicrobial resistance genes despite complete phenotypic susceptibility to the antimicrobial agents tested. The majority of identified determinants comprised chromosomally encoded multidrug efflux systems, regulatory networks, and intrinsic resistance-associated genes, including the AcrAB-TolC complex, AcrD, Mdt transporters, MarA, Emr systems, and the class C β-lactamases AmpC and AmpH (33,37,38). Nevertheless, the presence of these genes should not be interpreted as evidence of clinically expressed antimicrobial resistance because many constitute components of the intrinsic *E. coli* resistome whose contribution to resistance depends on transcriptional regulation, gene expression, membrane permeability, and environmental selective pressures (33,37–39). The concordance between phenotypic susceptibility and the absence of acquired high-impact resistance determinants indicates that the isolate remains susceptible under laboratory testing conditions. This finding emphasizes the importance of integrating phenotypic susceptibility testing with whole-genome sequencing when interpreting antimicrobial resistance profiles.

Plasmid analysis identified three Col-type plasmid replicons, suggesting the presence of mobile genetic elements that may facilitate horizontal gene transfer. Although the current short-read sequencing approach could not completely resolve plasmid architecture or definitively assign all genes to plasmids or the chromosome, the coexistence of plasmid replicons with the complete *bfp* operon is consistent with the established role of the EAF plasmid in typical EPEC pathogenesis (12,14). Long-read sequencing would provide improved resolution of plasmid structure and enable comprehensive characterization of mobile genetic elements associated with virulence and antimicrobial resistance.

Core genome phylogenetic analysis demonstrated that the bat-derived isolate (O128:H6) clustered within an established EPEC lineage and showed its closest relationship to the human-derived EPEC strain 1554 from the Netherlands. This finding indicates conservation of core genomic characteristics among geographically distinct EPEC lineages while illustrating the broad evolutionary diversity of the pathotype. Importantly, phylogenetic proximity alone should not be interpreted as evidence of recent transmission between wildlife and humans. Instead, these data demonstrate shared evolutionary ancestry among EPEC lineages and reinforce previous observations that pathogenic *E. coli* possess a highly conserved core genome upon which diverse combinations of virulence-associated mobile genetic elements are superimposed (6–8,53).

The detection of a genomically confirmed typical EPEC isolate in an Egyptian fruit bat contributes to the growing body of evidence that wildlife can harbor diarrheagenic *E. coli*. Previous investigations have detected EPEC-associated virulence genes in bats and other wildlife species, although relatively few studies have applied whole-genome sequencing to characterize complete virulence repertoires or evolutionary relationships (26,57). The present study therefore extends existing knowledge by demonstrating the complete genomic organization of a typical EPEC isolate recovered from a free-ranging bat in East Africa. Given the increasing overlap among wildlife, livestock, and human populations resulting from habitat modification, agricultural expansion, and urbanization, continued genomic surveillance of bacterial pathogens at wildlife–human interfaces is consistent with One Health strategies for identifying microorganisms of potential public health importance (21–26).

Identification of an isolate possessing virulence determinants characteristic of clinically important EPEC highlights the importance of including wildlife in integrated surveillance programs designed to monitor the distribution and evolution of diarrheagenic *E. coli*. Whole-genome sequencing provides an effective platform for detecting changes in virulence gene content, antimicrobial resistance determinants, and population structure that may precede the emergence of clinically significant lineages.

### Limitations[H1]

This study has several limitations. First, sampling was limited to a single geographic region and one sampling period, restricting inference regarding the wider distribution of EPEC among bat populations in Kenya. Second, only one genomically confirmed typical EPEC isolate was identified, limiting conclusions regarding prevalence or population structure. Third, short-read sequencing did not permit complete reconstruction of plasmids or definitive localization of all mobile genetic elements. Finally, functional assays evaluating adherence, gene expression, or pathogenicity were not performed, preventing assessment of the phenotypic activity of the identified virulence determinants.

### Conclusion

Whole-genome sequencing identified a typical EPEC isolate carrying the complete LEE pathogenicity island and bundle-forming pilus operon in a free-ranging Egyptian fruit bat from coastal Kenya. The isolate exhibited genomic characteristics closely resembling those of established human EPEC lineages while remaining phenotypically susceptible to all antimicrobial agents tested. These findings expand current understanding of the diversity of wildlife-associated diarrheagenic *E. coli* and underscore the value of genomic surveillance for monitoring the evolution of bacterial pathogens at wildlife–human interfaces within a One Health framework.

## Declarations

### Ethics approval and consent to participate

The study was approved by the Kenya Medical Research Institute (KEMRI) Scientific and Ethics Review Unit (SERU) under protocol number KEMRI/SERU/CCR/0353/4973, and also by the National Commission for Science, Technology & Innovation (NACOSTI) under license number 500230. Additional approval was done by the Animal Care and Use Review Office (ACURO), Walter Reed Army Institute of Research (WRAIR) under reference WRAIR 3187, with exemptions from further approvals granted as per WRAIR policy #25 since no samples, data or information was being collected from human subjects.

### Availability of data and material

The datasets supporting the conclusions made in this article are included within the document, and are also available from the corresponding author upon reasonable request. The sequence data generated are linked to the NCBI BioProject PRJNA1559230.

### Disclaimer

Material has been reviewed by the Walter Reed Army Institute of Research. There is no objection to its presentation and/or publication. The views expressed herein reflect the results of research conducted by the author(s) and do not necessarily reflect the official policy or position of the Defense Health Agency, the Department of War, the U.S. Government, nor the HJF Medical Research International, Inc,

### Author contribution

JNK: study and experimental design; bacteria culture; isolate identification; NGS sequencing; sequencing data analysis; manuscript development and writing, BK: NGS sequencing and sequencing data analysis; DA: sample collection, technical consultation, manuscript review; BA: sample collection, manuscript development; MWV: manuscript development; LM, EO, CKM, JO, HR, GGK, JE: technical consultation and manuscript review; FE: project conception; study and experimental design; data analysis; and manuscript development. The manuscript was read and approved by all the authors.

## References

1. Janda JM, Abbott SL. The changing face of the family Enterobacteriaceae (order: “Enterobacterales”): New members, taxonomic issues, geographic expansion, and new diseases and disease syndromes. Clin Microbiol Rev. 2021;34(2):e00174–20. doi:10.1128/CMR.00174-20

2. Croxen MA, Finlay BB. Molecular mechanisms of Escherichia coli pathogenicity. Nat Rev Microbiol. 2010 Jan 7;8(1):26–38. doi:10.1038/nrmicro2265

3. Manges AR, Geum HM, Guo A, Edens TJ, Fibke CD, Pitout JDD. Global Extraintestinal Pathogenic Escherichia coli (ExPEC) Lineages. Clin Microbiol Rev. 2019 Jun 19;32(3). doi:10.1128/CMR.00135-18

4. Kaper JB, Nataro JP, Mobley HL. Pathogenic Escherichia coli. Nat Rev Microbiol. 2004;2(2):123–40.

5. Bueris V, Sircili MP, Taddei CR, dos Santos MF, Franzolin MR, Martinez MB, et al. Detection of diarrheagenic Escherichia coli from children with and without diarrhea in Salvador, Bahia, Brazil. Mem Inst Oswaldo Cruz. 2007;102(7):839–44.

6. Rasko DA, Rosovitz MJ, Myers GSA, Mongodin EF, Fricke WF, Gajer P, et al. The pangenome structure of Escherichia coli: Comparative genomic analysis of E. coli commensal and pathogenic isolates. J Bacteriol. 2008 Oct;190(20):6881–93. doi:10.1128/JB.00619-08 PubMed PMID: 18676672.

7. Moriel DG, Rosini R, Seib KL, Serino L, Pizza M, Rappuoli R. Escherichia coli: Great diversity around a common core. mBio. 2012;3(3):e00118–12. doi:10.1128/mBio.00118-12

8. Croxen MA, Law RJ, Scholz R, Keeney KM, Wlodarska M, Finlay BB. Recent Advances in Understanding Enteric Pathogenic Escherichia coli. Clin Microbiol Rev. 2013 Oct;26(4):822–80. doi:10.1128/CMR.00022-13

9. Kotloff KL, Nataro JP, Blackwelder WC, Nasrin D, Farag TH, Panchalingam S, et al. Burden and aetiology of diarrhoeal disease in infants and young children in developing countries (the Global Enteric Multicenter Study, GEMS): A prospective, case-control study. The Lancet. 2013;382(9888):209–22.

10. Fagerli K, Omore R, Kim S, Ochieng JB, Ayers TL, Juma J, et al. Factors associated with typical enteropathogenic Escherichia coli infection among children <5 years old with moderate-to-severe diarrhoea in rural western Kenya, 2008–2012. Epidemiol Infect. 2020;148:e281. doi:10.1017/S0950268820002730

11. Ingle DJ, Tauschek M, Edwards DJ, Hocking DM, Pickard DJ, Azzopardi KI, et al. Evolution of atypical enteropathogenic E. coli by repeated acquisition of LEE pathogenicity island variants. Nat Microbiol. 2016;1(2):15010. doi:10.1038/nmicrobiol.2015.10

12. Epler Barbercheck CR, Bullitt E, Andersson M. Bacterial adhesion pili. In: Harris JR, Boekema EJ, editors. Membrane protein complexes: Structure and function. Springer; 2018. p. 1–18.

13. Cleary J, Lai LC, Shaw RK, Straatman-Iwanowska A, Donnenberg MS, Frankel G, et al. Enteropathogenic Escherichia coli (EPEC) adhesion to intestinal epithelial cells: Role of bundle-forming pili (BFP), EspA filaments and intimin. Microbiology (N Y). 2004;150(3):527–38.

14. Abe CM, Trabulsi LR, Blanco J, Blanco M, Dahbi G, Blanco JE, et al. Virulence features of atypical enteropathogenic Escherichia coli identified by the eae+ EAF-negative stx− genetic profile. Diagn Microbiol Infect Dis. 2009;64(4):357–65. doi:10.1016/j.diagmicrobio.2009.03.029

15. Yatsuyanagi J, Saito S, Miyajima Y, Amano K, Enomoto K. Characterization of atypical enteropathogenic Escherichia coli strains harboring the astA gene that were associated with a waterborne outbreak of diarrhea in Japan. J Clin Microbiol. 2003;41(5):2033–9.

16. Brzuszkiewicz E, Thürmer A, Schuldes J, Leimbach A, Liesegang H, Meyer FD, et al. Genome sequence analyses of two isolates from the recent Escherichia coli outbreak in Germany reveal the emergence of a new pathotype: Entero-aggregative-haemorrhagic Escherichia coli (EAHEC). Arch Microbiol. 2011;193(12):883–91. doi:10.1007/s00203-011-0725-6

17. Trabulsi LR, Keller R, Gomes TAT. Typical and Atypical Enteropathogenic Escherichia coli. Emerging Infectious Diseases •. 2002.

18. Nataro JP, Kaper JB. Diarrheagenic Escherichia coli [Internet]. Vol. 11. 1998. Available from: https://journals.asm.org/journal/cmr

19. Farina LL, Lankton JS. Chiroptera. In: Pathology of Wildlife and Zoo Animals. Elsevier; 2018. p. 607–33. doi:10.1016/B978-0-12-805306-5.00025-0

20. Fleming TH. Bat Migration. In: Encyclopedia of Animal Behavior, Second Edition: Volume 1-5. Elsevier; 2019. p. V3-605-V3-610. doi:10.1016/B978-0-12-809633-8.20764-4

21. Kuzmin I V., Bozick B, Guagliardo SA, Kunkel R, Shak JR, Tong S, et al. Bats, emerging infectious diseases, and the rabies paradigm revisited. Emerging Health Threats Journal. 2011. doi:10.3402/ehtj.v4i0.7159

22. Brook CE, Dobson AP. Bats as ‘special’ reservoirs for emerging zoonotic pathogens. Trends in Microbiology. Elsevier Ltd; 2015. p. 172–80. doi:10.1016/j.tim.2014.12.004 PubMed PMID: 25572882.

23. Allocati N, Petrucci AG, Di Giovanni P, Masulli M, Di Ilio C, De Laurenzi V. Bat–man disease transmission: zoonotic pathogens from wildlife reservoirs to human populations. Cell Death Discovery. Springer Nature; 2016. doi:10.1038/cddiscovery.2016.48 PubMed PMID: 27551536.

24. Hayman DTS. Bats as Viral Reservoirs. Annual Review of Virology. Annual Reviews Inc.; 2016. p. 77–99. doi:10.1146/annurev-virology-110615-042203 PubMed PMID: 27578437.

25. Calisher CH, Childs JE, Field HE, Holmes K V., Schountz T. Bats: Important reservoir hosts of emerging viruses. Clinical Microbiology Reviews. 2006. p. 531–45. doi:10.1128/CMR.00017-06 PubMed PMID: 16847084.

26. Mühldorfer K. Bats and Bacterial Pathogens: A Review. Zoonoses Public Health. 2013 Feb;60(1):93–103. doi:10.1111/j.1863-2378.2012.01536.x PubMed PMID: 22862791.

27. Barry A. An Overview of the Clinical and Laboratory Standards Institute (CLSI) and Its Impact on Antimicrobial Susceptibility Tests. In: Antimicrobial Susceptibility Testing Protocols. CRC Press; 2007. p. 1–6. doi:10.1201/9781420014495.ch1

28. Shen Z. DNA Extraction with Zymo Quick-DNATM Fungal/Bacterial Miniprep Kit v1. 2025. doi:10.17504/protocols.io.dm6gpdw98gzp/v1

29. Habets A, Touzain F, Lucas P, Huong NTT, Iguchi A, Crombé F, et al. Identification of Five Serotypes of Enteropathogenic Escherichia coli from Diarrheic Calves and Healthy Cattle in Belgium and Comparative Genomics with Shigatoxigenic E. coli. Vet Sci. 2022 Sep 1;9(9). doi:10.3390/vetsci9090492

30. Blanco M, Schumacher S, Tasara T, Zweifel C, Blanco JE, Dahbi G, et al. Serotypes, intimin variants and other virulence factors of eae positive Escherichia coli strains isolated from healthy cattle in Switzerland. Identification of a new intimin variant gene (eae-η2). BMC Microbiol. 2005 May 9;5. doi:10.1186/1471-2180-5-23 PubMed PMID: 15882459.

31. Higgins J, Hohn C, Hornor S, Frana M, Denver M, Joerger R. Genotyping of Escherichia coli from environmental and animal samples. J Microbiol Methods. 2007;70(2):227–35.

32. Ishii S, Meyer KP, Sadowsky MJ. Relationship between phylogenetic groups, genotypic clusters, and virulence gene profiles of Escherichia coli strains from diverse human and animal sources. Appl Environ Microbiol. 2007 Sep;73(18):5703–10. doi:10.1128/AEM.00275-07 PubMed PMID: 17644637.

33. Fernando D, Kumar A. Resistance-Nodulation-Division Multidrug Efflux Pumps in Gram-Negative Bacteria: Role in Virulence. Antibiotics. 2013 Mar 18;2(1):163–81. doi:10.3390/antibiotics2010163

34. Ghachi M El, Bouhss A, Blanot D, Mengin-Lecreulx D. The bacA Gene of Escherichia coli Encodes an Undecaprenyl Pyrophosphate Phosphatase Activity. Journal of Biological Chemistry. 2004 Jul;279(29):30106–13. doi:10.1074/jbc.M401701200

35. Sasal WM, Gmiter D, Kaca W. Polymyxin resistance in Gram-negative bacteria: a comprehensive review with a focus on the role of 4-amino-4-doexy-l-arabinose. Mol Biol Rep. 2026 Dec 29;53(1):223. doi:10.1007/s11033-025-11368-4

36. Moussatova A, Kandt C, O’Mara ML, Tieleman DP. ATP-binding cassette transporters in Escherichia coli. Biochimica et Biophysica Acta (BBA) - Biomembranes. 2008 Sep;1778(9):1757–71. doi:10.1016/j.bbamem.2008.06.009

37. Pao SS, Paulsen IT, Saier MH. Major Facilitator Superfamily. Microbiology and Molecular Biology Reviews. 1998 Mar;62(1):1–34. doi:10.1128/MMBR.62.1.1-34.1998

38. Philippon A, Arlet G, Labia R, Iorga BI. Class C β-Lactamases: Molecular Characteristics. Clin Microbiol Rev. 2022 Sep 21;35(3). doi:10.1128/cmr.00150-21

39. Ueguchi C, Mizuno T. The Escherichia coli nucleoid protein H-NS functions directly as a transcriptional repressor. EMBO J. 1993 Mar;12(3):1039–46. doi:10.1002/j.1460-2075.1993.tb05745.x

40. Dorman CJ, Higgins CF. Fimbrial phase variation in Escherichia coli: dependence on integration host factor and homologies with other site-specific recombinases. J Bacteriol. 1987 Aug;169(8):3840–3. doi:10.1128/jb.169.8.3840-3843.1987

41. Iguchi A, Thomson NR, Ogura Y, Saunders D, Ooka T, Henderson IR, et al. Complete genome sequence and comparative genome analysis of enteropathogenic Escherichia coli O127:H6 strain E2348/69. J Bacteriol. 2009 Jan;91(1):347–54. doi:10.1128/JB.01238-08 PubMed PMID: 18952797.

42. Lim JY, Yoon J, Hovde CJ. A brief overview of Escherichia coli O157:H7 and its plasmid O157. J Microbiol Biotechnol. 2010 Jan;20(1):5–14. PubMed PMID: 20134227.

43. Ochoa TJ, Salazar-Lindo E, Cleary TG. Evaluation of children with persistent infectious diarrhea. Semin Pediatr Infect Dis. 2004;16(4):259–63.

44. Abba K, Sinfield R, Hart CA, Garner P. Pathogens associated with persistent diarrhoea in children in low and middle income countries: Systematic review. BMC Infect Dis. 2009;9(88). doi:10.1186/1471-2334-9-88

45. Donnenberg MS, Hazen TH, Rasko DA. Bacterial factors associated with lethal outcome of enteropathogenic Escherichia coli infection: Genomic case-control studies. PLoS Negl Trop Dis. 2015;9(5):e0003791. doi:10.1371/journal.pntd.0003791

46. Kaper JB, Nataro JP, Mobley HLT. Pathogenic Escherichia coli. Nat Rev Microbiol. 2004 Feb;2(2):123–40. doi:10.1038/nrmicro818

47. Schmidt J, Zdarska V, Kolar M, Mlynarcik P. Analysis of BlaEC family class C beta-lactamase. FEMS Microbiol Lett. 2023;370. doi:10.1093/femsle/fnad097 PubMed PMID: 37757475.

48. Paterson DL, Bonomo RA. Extended-Spectrum β-Lactamases: a Clinical Update. Clin Microbiol Rev. 2005 Oct;18(4):657–86. doi:10.1128/CMR.18.4.657-686.2005

49. Cantón R, Coque TM. The CTX-M β-lactamase pandemic. Curr Opin Microbiol. 2006 Oct;9(5):466–75. doi:10.1016/j.mib.2006.08.011

50. Livermore DM. Fourteen years in resistance. Int J Antimicrob Agents. 2012 Apr;39(4):283–94. doi:10.1016/j.ijantimicag.2011.12.012

51. Gyansa-Lutterodt M. Antibiotic resistance in Ghana. Lancet Infect Dis. 2013 Dec;13(12):1006–7. doi:10.1016/S1473-3099(13)70196-8

52. Laxminarayan R, Duse A, Wattal C, Zaidi AKM, Wertheim HFL, Sumpradit N, et al. Antibiotic resistance—the need for global solutions. Lancet Infect Dis. 2013 Dec;13(12):1057–98. doi:10.1016/S1473-3099(13)70318-9

53. Tenaillon O, Skurnik D, Picard B, Denamur E. The population genetics of commensal Escherichia coli. Nat Rev Microbiol. 2010 Mar;8(3):207–17. doi:10.1038/nrmicro2298

54. Deborah Chen H, Frankel G. Enteropathogenic Escherichia coli: Unravelling pathogenesis. FEMS Microbiology Reviews. 2005. p. 83–98. doi:10.1016/j.femsre.2004.07.002 PubMed PMID: 15652977.

55. Ochoa TJ, Barletta F, Contreras C, Mercado E. New insights into the epidemiology of enteropathogenic Escherichia coli infection. Transactions of the Royal Society of Tropical Medicine and Hygiene. 2008. p. 852–6. doi:10.1016/j.trstmh.2008.03.017 PubMed PMID: 18455741.

56. African Fruit Bats as Potential Reservoir for Zoonotic Pathogens - The Example of Escherichia coli.

57. Cabal A, Pereira MJ, Aguiar LMS, Domínguez L, Fonseca C, Álvarez J, et al. Direct detection of Escherichia coli virulence genes by real-time PCR in fecal samples from bats in Brazil. J Wildl Dis. 2015 Oct 1;51(4):942–5. doi:10.7589/2015-01-024 PubMed PMID: 26267461.

